# IFN-α Induces Heterogenous ROS Production in Human β-Cells

**DOI:** 10.1101/2025.02.19.639120

**Authors:** Leslie E. Wagner, Olha Melnyk, Abigail Turner, Bryce E. Duffett, Charanya Muralidharan, Michelle M. Martinez-Irizarry, Matthew C. Arvin, Kara S. Orr, Elisabetta Manduchi, Klaus H. Kaestner, Joseph T. Brozinick, Amelia K. Linnemann

**Affiliations:** Departments of Biochemistry and Molecular Biology, Indiana University School of Medicine, Indianapolis, IN; Departments of Pediatrics, Indiana University School of Medicine, Indianapolis, IN; Departments of Center for Diabetes and Metabolic Diseases, Indiana University School of Medicine, Indianapolis, IN; Department of Genetics and Institute for Diabetes, Obesity and Metabolism, University of Pennsylvania, Philadelphia, PA; Eli Lilly and Company, Indianapolis, IN

## Abstract

Type 1 diabetes (T1D) is a multifactorial disease involving genetic and environmental factors, including viral infection. We investigated the impact of interferon alpha (IFN-α), a cytokine produced during the immune response to viral infection or the presence of un-edited endogenous double-stranded RNAs, on human β-cell physiology. Intravital microscopy on transplanted human islets using a β-cell-selective reactive oxygen species (ROS) biosensor (RIP1-GRX1-roGFP2), revealed a subset of human β-cells that acutely produce ROS in response to IFN-α. Comparison to Integrated Islet Distribution Program (IIDP) phenotypic data revealed that healthier donors had more ROS accumulating cells. I*n vitro* IFN-α treatment of human islets similarly elicited a heterogenous increase in superoxide production that originated in the mitochondria. To determine the unique molecular signature predisposing cells to IFN-α stimulated ROS production, we flow sorted human islets treated with IFN-α. RNA sequencing identified genes involved in inflammatory and immune response in the ROS-producing cells. Comparison with single cell RNA-Seq datasets available through the Human Pancreas Analysis Program (HPAP) showed that genes upregulated in ROS-producing cells are enriched in control β-cells rather than T1D donors. Combined, these data suggest that IFN-α stimulates mitochondrial ROS production in healthy human β-cells, potentially predicting a more efficient antiviral response.

## INTRODUCTION

Type 1 diabetes (T1D) has long been described as a multifactorial disease, consisting of both a genetic predisposition and an environmental triggering event. A leading hypothesis for this environmental trigger is early childhood viral infection leading to viral mimicry and spread of T cell immunity to beta-cell antigens. Interferon alpha (IFN-α), a type I interferon (IFN), is a pluripotent inflammatory cytokine produced by cells involved in the innate immune response during viral infection, or in response to unedited endogenous double stranded RNAs. IFN-α and other type 1 interferons likely play a role in T1D pathophysiology. For example, children with a genetic predisposition for T1D development have a type I interferon-inducible transcriptional signature in their serum preceding the detection of autoantibodies (1, 2). Islets isolated from newly diagnosed donors have been shown to have both IFN-α expression and an increase in type 1 IFN-stimulated genes (3–5). Additionally, studies conducted with non-diabetic human islets have shown that IFN-α alone induces MHC-I overexpression and ER stress, two hallmarks of early T1D, further suggesting its role in disease development (6).

We and others have shown that proinflammatory cytokines increase reactive oxygen species (ROS) in β-cells (7–10). ROS are chemically reactive, unstable compounds derived from oxygen. Most ROS are naturally produced in β-cells as byproducts of metabolic reactions and can stimulate important cellular processes such as β-cell proliferation (11) and glucose stimulated insulin secretion (12, 13). However, when the levels of ROS increase above what can be detoxified by scavenging enzymes in the antioxidant response, oxidative stress ensues and can lead to cellular dysfunction and destruction (14–16).

In recent years, β-cell heterogeneity has been a well-explored topic, with β-cells displaying heterogeneity with both gene expression (17–20) and cellular function to include functional heterogeneity in insulin secretion (21) and calcium signaling (22–24), suggesting that not all β-cells respond to stimuli in the same manner. Additionally, although T1D is an autoimmune disorder characterized by immune-mediated destruction of the insulin-producing β-cells in the pancreatic islet, recent evidence shows residual insulin-positive β-cells in the islets of people with longstanding T1D, suggesting cell survival heterogeneity also exists in this population (25–27). The nonobese diabetic (NOD) mouse model of autoimmune diabetes has also been studied in this context, and a similar subpopulation of persistent β-cells resisting sustained immune attack has been identified (28). Together, this suggests that a small subpopulation of uncharacterized β-cells exists that are innately resistant to immune assault.

Utilizing intravital microscopy, we show here evidence that following IFN-α stimulation, human β-cells undergo an acute heterogenous increase in ROS production *in vivo*. Interestingly, not all donors displayed this response, with some donors eliciting no β-cell ROS production in response to IFN-α. A review of donor phenotypic information revealed that healthier donors, i.e., those with lower BMI and HbA1c levels, had the highest number of cells that accumulate ROS. Analysis of up-regulated genes in ROS accumulating cells also demonstrated an enrichment in genes whose β-cell expression is higher in controls than in T1D donors. Our data therefore suggests that ROS accumulation in response to acute type 1 interferon stress may be protective and could play an important role in the β-cell’s adaptive response to viral infection.

## RESULTS

### IFN-α Induces Heterogenous ROS Accumulation in Human Islets *in vivo*

Human islets were received from the Integrated Islet Distribution Program (IIDP) or Alberta Diabetes Institute (ADI) and transfected with 1*10^11^ PFU/1,000 IEQs of adenoviral RIP1-Grx1-roGFP2, a previously established, ratiometric ROS biosensor that we placed under the insulin promoter (29). See Table 1 for donor details. In the presence of ROS in the cytoplasm, there is a conformational change in GFP, resulting in the sensor being more excitable at 405nm. Therefore, cells undergoing ROS production can be identified by an increase in the 405nm signal and decrease in the 488nm signal (Fig 1A). This expected change in the biosensor excitation is shown in Fig 1B. Two weeks following transplantation of human islets, the kidney containing transduced islets was externalized on an imaging dish for real-time monitoring of the biosensor response. At this timepoint, we observed robust vascularization of the engrafted islets (Supplementary Fig. 1A) along with the presence of human C-peptide in all animals (Supplementary Fig. 1B). Following baseline image acquisition, 2.75*10^5^ IU/mL of carrier-free recombinant human IFN-α or saline was retro-orbitally administered. The biosensor response was measured for 60 minutes following treatment injection and cells were individually traced to define regions of interest (ROIs) for analysis and followed throughout the time of imaging.

**Table 1:** Human Islet Donors Used for Experiments. Phenotypic information from each donor, including sex, age, BMI, and HbA1C are provided.

| Donor ID | Source | Sex | Age | BMI | HbA1C | Experiment |
| --- | --- | --- | --- | --- | --- | --- |
| SAMN10920290 | IIDP | F | 46 | 18.2 | 4.4 | Intravital imaging |
| R309 | Alberta | F | 47 | 27.4 | 5.5 | Intravital imaging |
| SAMN17762820 (R395) | Alberta | F | 28 | 31.4 | 3.6 | Intravital imaging |
| SAMN16427178 | IIDP | F | 42 | 31.2 | 5.5 | Intravital imaging |
| SAMN16423389 | IIDP | M | 55 | 25.8 | 4.9 | Intravital imaging |
| SAMN17367763 | IIDP | M | 56 | 33 | 5.3 | Intravital imaging |
| SAMN28450743 | IIDP | M | 47 | 25.5 | 5.1 | Intravital imaging |
| SAMN29094695 (R448) | Alberta | F | 61 | 36.1 | 5.8 | Intravital imaging |
| SAMN38094145 | IIDP | F | 52 | 33.6 | 5.7 | DHE *in vitro* imaging |
| SAMN37217050 (R505) | Alberta | F | 53 | 33.1 | 5.4 | DHE *in vitro* imaging |
| SAMN37844676 (R511) | Alberta | F | 47 | 19.2 | 5.9 | DHE *in vitro* imaging |
| SAMN37844810 (R512) | Alberta | M | 63 | 34.7 | 5.9 | DHE *in vitro* imaging |
| SAMN38760239 (R522) | Alberta | M | 56 | 29.4 | 5 | DHE *in vitro* imaging |
| SAMN40137906 (R532) | Alberta | F | 28 | 24.3 | 5.5 | DCFDA *in vitro* imaging |
| SAMN40723343 (R538) | Alberta | M | 61 | 32.9 | 5.5 | DCFDA *in vitro* imaging |
| SAMN40709610 | IIDP | F | 55 | 28.5 | 5.7 | DCFDA *in vitro* imaging |
| R506 (SAMN37217101) | Alberta | M | 66 | 32.4 | 7.6 | DHE *in vitro* imaging- T2D |
| SAMN40974624 | IIDP | M | 44 | 34.8 | 6.5 | DHE *in vitro* imaging- T2D |
| SAMN41264680 | IIDP | M | 52 | 27.8 | 8.5 | DHE *in vitro* imaging- T2D |
| SAMN38891395 | IIDP | F | 71 | 29 | 5.1 | RNA Sequencing |
| SAMN38872001 | IIDP | F | 54 | 39.7 | 5.9 | RNA Sequencing |
| SAMN39461369 | IIDP | F | 53 | 24.7 | 5.6 | RNA Sequencing |
| SAMN35985448 (R496) | Alberta | F | 68 | 22.3 | 5.6 | Immunofluorescence staining |

**Figure 1:**
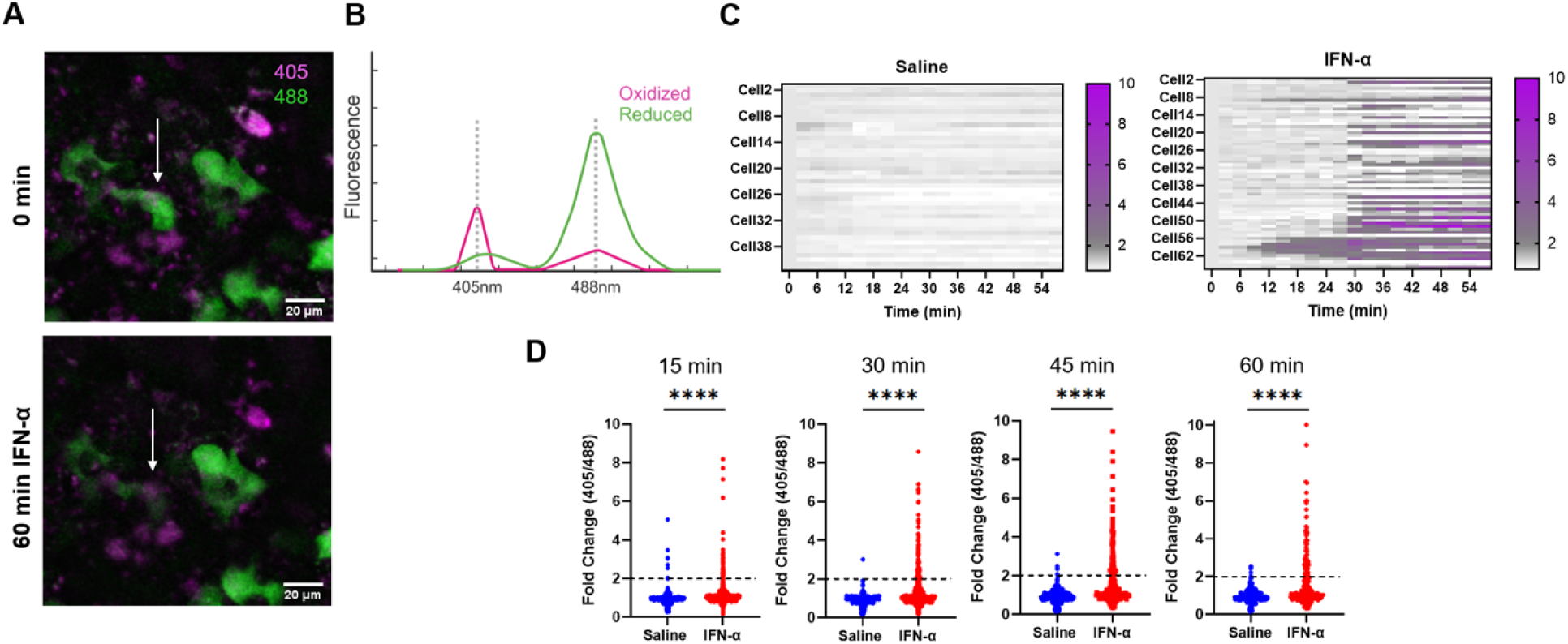
Human β-cells Accumulate ROS *in vivo* Following IFN-α Stimulation. (A) Representative images of a responding β-cell at baseline and 60 minutes following IFN-α stimulation (white arrow). Using the RIP1-GRX1-roGFP2 biosensor, responding β-cells can be identified with an increase in 405nm channel intensity (magenta) and a decrease in 488nm channel intensity (green). (B) In the presence of ROS, conformational changes in the RIP1-GRX1-roGFP2 biosensor make it more exciteable at 405nm, allowing the detection of oxidized cells. (C) Representative heatmaps from donors with islets stimulated with saline or IFN-α. Plotted in the heatmaps is the fold change in 405nm/488nm ratio, relative to baseline, of the biosensor for single cells during the entire course of imaging. Cells that have a fold change of 2 or higher are represented in magenta. (D) Graphs depicting the fold change in single cells from all donors combined at 15 min, 30 min, 45 min, and 60 min. Saline stimulated cells are depicted in blue, and IFN-α stimulated cells are depicted in red. A dotted line is placed marking a fold change of 2. Analysis was conducted as a t-test, comparing the means of both treatments.

Approximately 30 minutes after IFN-α administration, we observed that a subset (0-63%) of human β-cells exhibit a robust production of ROS, as measured by the fold change in biosensor ratio (405nm/488nm) compared to baseline (Fig 1C), with some of these ‘responder’ cells accumulating ROS as early as 15 minutes after IFN-α administration (Fig. 1C). Across multiple donors, we observe that β-cell ROS accumulation peaks at around 45 min following IFN-α stimulation (Fig 1D). All animals had measurable circulating IFN-α levels at the end of the imaging session (Supplementary Fig. 1C).

### Donors with Higher Proportion of Responders Are Healthier

We observed a significant amount of heterogeneity between donors, where roughly half of donors studied did not produce significant amounts of ROS in response to IFN-α production (Fig 2A). Therefore, we asked if this response was due to phenotypic differences among donors. Using the Human Islet Phenotyping Program data provided by IIDP, we integrated our data with phenotypic information including sex, age, BMI, HbA1c, ethnicity/race, perifusion data, islet insulin content, and endocrine cell composition. Plotting these variables against % responders from each donor at the 30-minute timepoint, we identified a strong negative correlation between percent responders with both BMI and HbA1c (Fig 2B and Fig 2C). We did not observe strong correlations with any of the other variables provided (Supplementary Fig 2). Our observation that ‘healthier’ donors had a higher number of cells with increased ROS in response to IFN-α suggest that this response may be a beneficial adaptive stress response to external cellular stress such as cytokines.

**Figure 2:**
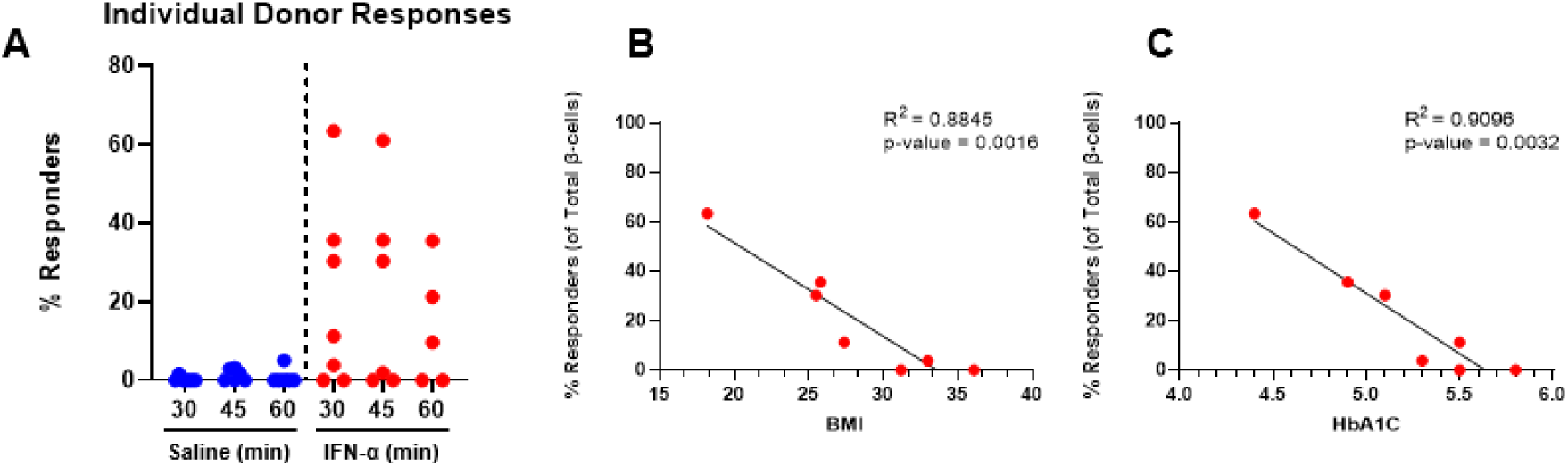
Healthier Donors Have a Higher Proportion of Responders. (A) The percentage of responders from total β-cells are plotted for each donor used in intravital imaging experiments at 30, 45, and 60 min timepoints. The percentage of saline responders are shown in blue, and the percentage of IFN-α responders are shown in red. (B) The BMI of each donor is plotted against the donor’s percent responders at 30 min. A linear regression analysis was conducted, and the R^2^ and p-values from this analysis are shown. (C) The HbA1C of each donor is plotted against the donor’s percent responders at 30 min. A linear regression analysis was conducted, and the R^2^ and p-values from this analysis are shown.

### IFN-α Treatment Induces Acute Protein Changes in IFN-α Signaling and ROS Production

To validate that β-cells also exhibit evidence of expected signaling downstream of the IFN-α receptor, kidneys containing transplanted islets were embedded and further analyzed. We observed a broad increase in nuclear pSTAT2 expression, a transcription factor in the IFN-α signaling pathway, in islets stimulated with IFN-α compared to those from mice receiving saline (Supplementary Fig. 3A), suggesting that there is no difference in STAT-mediated IFN-α signaling between β-cells. This is consistent with the species specificity of cytokines expected with injection of human IFN-α as well as our observation of human IFN-α only in the circulation of the IFN-α injected animals (Supplementary Fig. 1C). Additionally, islet transplants were stained for 8-Hydroxy-2’-deoxyguanosine (8OHdG) and 4-hydroxy-2-nonenal (4-HNE), markers of oxidative stress. We observed a heterogenous nuclear increase in both 8OHdG (Supplementary Fig. 3B) and 4-HNE (Supplementary Fig 3C) signal in IFN-α stimulated islets that was absent in saline stimulated islets. These data demonstrate that the IFN-α likely reaches all but promotes the generation of ROS in only a subset of β-cells.

### IFN-α Induces Heterogenous ROS Accumulation in Human Islets *in vitro*

Once we determined that IFN-α stimulates ROS production in a subset of human β-cells, we next wanted to identify the specific ROS being produced to elucidate a potential mechanism. Human islets were obtained from either the IIDP or the ADI, as above, and were rested in an incubator overnight (Table 1). Following treatment with 2,000 IU/mL recombinant human IFN-α and appropriate controls for 1 hour, islets were placed in fresh media containing 2μM Hoechst and 10μΜ dihydroethidium (DHE) or 5μM 2’,7’-dichlorodihydrofluorescein diacetate (DCFDA) for detection of superoxide or hydrogen peroxide production, respectively. In the presence of superoxide, a conformational change in DHE occurs, allowing it to intercalate with nucleic acids, resulting in a strong, nuclear stain(30). In the presence of hydrogen peroxide, DCFDA undergoes a conformational change, allowing it to become fluorescent(31). As demonstrated in Fig 3A, IFN-α induces a heterogenous accumulation of superoxide from healthy control donors *in vitro,* demonstrated by a non-uniform increase in nuclear DHE signal. Overall, we found that IFN-α significantly increased the density of DHE positive nuclei, as well as the percentage of DHE positive nuclei (Fig 3B). In contrast, we observed no statistically significant changes in DCFDA intensity (Fig 3B), suggesting that hydrogen peroxide is not a major factor in this process. When treating islets isolated from type 2 diabetic (T2D) donors with the same conditions *in vitro*, this response is greatly blunted (Fig 3A), as shown by a decreased occurrence and intensity of nuclear DHE staining, even in the presence of a positive control, Antimycin A(13). Quantitative analysis revealed that IFN-α does not stimulate an increase in the number of DHE positive nuclei per μm^2^ or an increase in the total percentage of positive nuclei in islets from T2D donors (Fig 3C).

**Figure 3:**
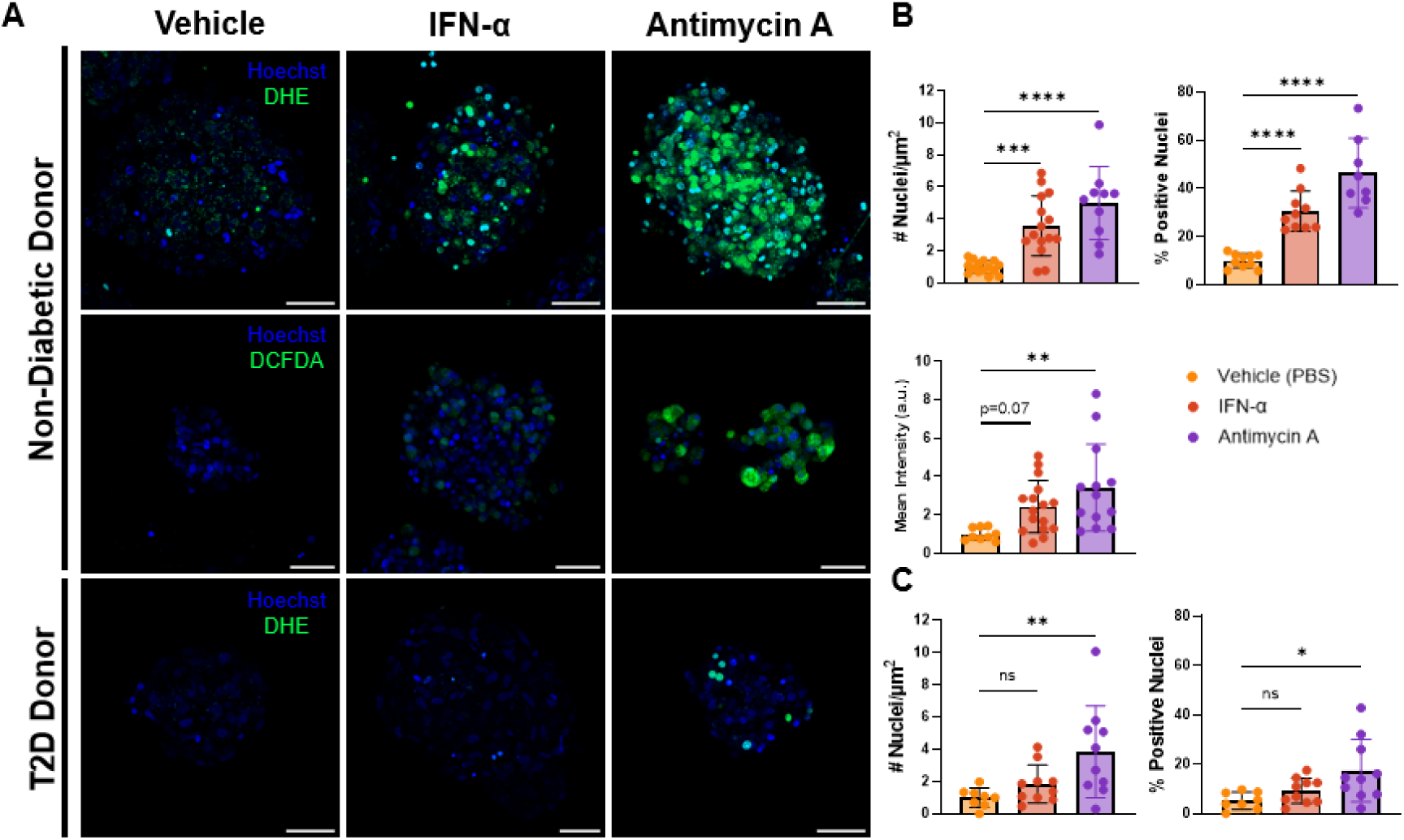
Human Islets Accumulate ROS *in vitro* Following IFN-α Stimulation. Human islets were treated *in vitro* with IFN-α or cooresponding controls. PBS was used for the vehicle condition, and Antimycin A was used for the positive control. Following treatment, human islets were stained with Hoechst (blue) for identification of nuclei and either DHE or DCFDA (green) for measurement of ROS. (A) Representative images of human islets stained with Hoechst and ROS probes from non-diabetic and type 2 diabetic donors. Images are maximum intensity projections of a Z-stack image collected throughout the islet. Scale bar length is 50μm. (B) Quanitfication of the non-diabetic donor images for both DHE and DCFDA. For DHE, images were quantified as the number of positive nuclei per μm^2^ of islet area and the percentage of DHE-positive nuclei. Analysis is represented as a fold change from the vehicle condition for each day a replicate was conducted. A one-way ANOVA test was conducted, with variation plotted as the mean±SD. For DCFDA analysis, the mean intensity of DCFDA was plotted and analyzed by a one-way ANOVA test, with variation plotted as mean±SD. The * denotes a p-value less than 0.05, ** denotes a p-value less than 0.01, *** denotes a p-value less than 0.001, and the **** denotes a p-value less than 0.0001. (C) Quanitfication of the diabetic donor images with DHE staining. Analysis was conducted as described in (B).

### IFN-α Induces an Inflammatory Transcriptional Response in Cells Producing ROS

To identify transcriptional network differences that might predispose a specific population of islet cells to ROS accumulation in response to IFN-α, we acutely treated human islets from three human islet donors with vehicle and 2,000 IU/mL IFN-α for 1 hour as above. Following treatment, we dissociated islets, stained the cells with DHE, and flow sorted the populations both negative and positive for ROS based upon DHE fluorescence intensity, using Antimycin A as a positive control for gating (Fig 4A and Fig 5A). We then isolated RNA from these populations and conducted ultralow-mRNA sequencing on these samples. First, we compared the differences between the vehicle treated cells exhibiting basal ROS production and the responder population to identify unique effects of IFN-α stimulation on ROS-producing cells. We identified 296 upregulated and 386 downregulated genes from this analysis (Fig 4B). We conducted gene ontology (GO) analysis on these differentially expressed genes and identified several pathways upregulated in the IFN-α responder population as compared to cells that produce ROS at baseline, including several antiviral defense pathways such as response to virus, defense response to virus, positive regulation of MDA-5 signaling, cellular response to xenobiotic stress, positive regulation of RIG-I signaling, innate immune response, and inflammatory response (Fig 4C). Of note, several of these genes are involved in the MDA5 and RIG-I pathways to combat viral infection or respond to insufficiently edited endogenous dsRNAs, including both RIG-I (*DDX58* (adj. p-value: 6.075×10^−11^)) and MDA5 (*IFIH1* (adj. p-value: 2.026×10^−7^)) themselves.

**Figure 4:**
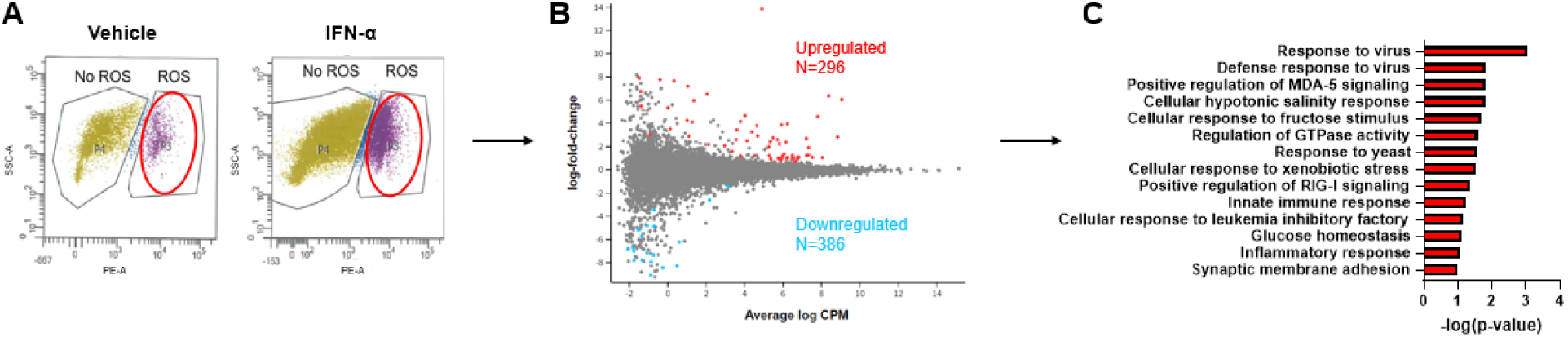
IFN-α Induces an Antiviral Transcriptional Response in Cells Producing ROS. Human islets were treated with vehicle or 2000 IU/mL IFN-α, dissociated, and stained with 10μM DHE to detect ROS. Samples were sorted based off the presence or absence of reactive oxygen species (ROS) as determined by established gates. (A) Representative image of how treated cells were sorted into ROS negative and ROS positive populations and collected for subsequent analysis. (B) Glimma plot of differentially expressed genes between the vehicle ROS positive and IFN-α ROS positive populations. Significantly upregulated genes are shown in red, and significantly downregulated genes are shown in cyan. (C) Significantly upregulated pathways in the IFN-α ROS positive (responder) population when compared to the vehicle ROS positive population, as determined with GO analysis. Values are represented as the –log(p-value) for each pathway.

**Figure 5:**
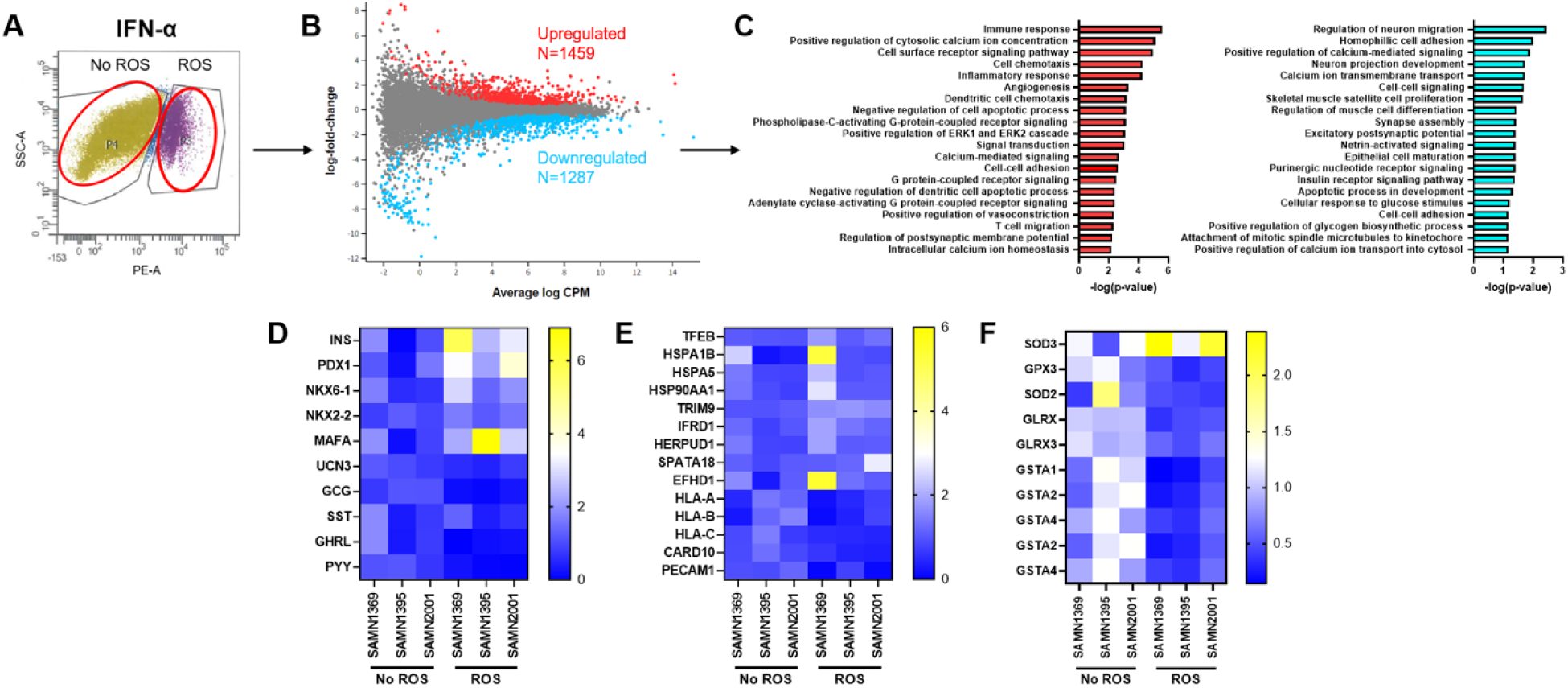
Cells Accumulating ROS in Response to IFN-α Have a Unique Molecular Signature. Human islets were treated with 2000 IU/mL IFN-α, dissociated, and stained with 10μM DHE to detect ROS. Samples were sorted based off the presence or absence of reactive oxygen species (ROS) as determined by established gates. (A) Representative image of how islet cells were sorted into ROS negative and ROS positive populations and collected for subsequent analysis. (B) Glimma plot of differentially expressed genes between the IFN-α ROS negative and IFN-α ROS positive populations. Significantly upregulated genes are shown in red, and significantly downregulated genes are shown in cyan. (C) Significantly upregulated (red) and downregulated (cyan) pathways in the IFN-α ROS positive (responder) population when compared to the IFN-α ROS negative population, as determined with GO analysis. Values are represented as the –log(p-value) for each pathway. (D) Heatmap detailing the expression of various islet cell markers. Heatmaps represent the fold change of gene expression compared to the vehicle negative for ROS population. Yellow represents more highly expressed genes, and blue denotes less expressed genes. (E) Heatmap detailing the expression of various antioxidant enzymes. (F) Heatmap detailing the expression of various cell survival and stress response markers.

### Cells Accumulating ROS in Response to IFN-α Have a Unique Molecular Signature

Once we determined the unique effect of IFN-α stimulation on ROS-producing human islet cells, we next wanted to identify differences in gene expression of human islet cells that produce ROS in response to IFN-α stimulation from those that do not, even when exposed to IFN-α. Therefore, we compared differentially expressed genes from the IFN-α treated negative for ROS population (termed ‘non-responders’) and the IFN-α responder population. Since the treatment time was only 1 hour, we rationalized that most of the differentially expressed genes (aside from the immediate response genes described in Fig. 4) would represent a signature that predisposes cells to ROS accumulation. Our analysis revealed 1,459 upregulated genes and 1,287 downregulated genes in the responder population (Fig 5B). Importantly, neither of the IFN-α receptors (*IFNAR1* and *IFNAR2*) were found to be differentially expressed between these two populations (Supp Fig 4). Ontology analysis of the differentially expressed genes revealed several upregulated pathways in the responder population, including immune response, inflammatory response, negative regulation of apoptosis, and modulation of calcium signaling (Fig 5C). Since we conducted this analysis on whole human islets and DHE is not β-cell-selective, we next determined if the responding population was enriched in any specific islet cell types. We found that responders had a higher expression of several β-cell identity markers, such as *INS*, *PDX1*, and *NKX2-2*, suggesting that this population is enriched for β-cells (Fig 5D). Additionally, the responder cells had a significantly lower expression of *GCG*, an α-cell marker, suggesting that the responders also contained a lower number of α-cells when compared to non-responders (Fig 5D).

Since ROS levels are mitigated in the cell by the antioxidant response, we next investigated the expression levels of several antioxidants between the non-responder and responder populations (Fig 5E). Interestingly, the expression of several genes encoding antioxidant-producing enzymes was decreased in the responder population, including cytoplasmic antioxidant enzymes (*GPX3*, *GST*, and *GLRX)* and the mitochondrial antioxidant enzyme *SOD2*. Additionally, since our results at this point suggested that ROS accumulation in response to IFN-α might be beneficial to the cell, we asked if responders exhibited any difference in genes involved in cell survival and stress response pathways. We observed increased expression of several pro-survival and stress response genes, including several genes involved in the cellular stress response to viral pathogens (Fig. 5F). This included several upregulated genes involved in the management of endoplasmic reticulum stress (*HSPA1B*, *HSPA5*, *HSP90AA1*, *HERPUD1*, and *TFEB*), regulation of cytokine production and inflammation (*IFRD1* and *CARD10*), and mitochondrial quality control (*SPATA18* and *EFHD1*). Additionally, responders have a decreased gene expression of *HLA-A*, *HLA-B*, and *HLA-C*, genes encoding HLA-I molecules that are implicated in the T-cell mediated destruction of beta cells in T1D pathogenesis.

### Responders are Enriched in Genes Negatively Correlated with T1D

To investigate if there was any relationship between differentially expressed genes in the IFN-α responder population and gene expression in the T1D disease state, we compared the genes differentially expressed in ROS accumulating IFN-α treated cells with single cell sequencing data collected by the Human Pancreas Analysis Program(32, 33) (HPAP; https://hpap.pmacs.upenn.edu). Gene Set Enrichment Analysis (GSEA; (34)) of islet scRNA_Seq data from donors with T1D and respective control donors without diabetes revealed that genes up-regulated in the IFN-α responder population were significantly enriched in genes with higher beta cell expression in the non-diabetic controls (Figs. 6A). Gene ontology analysis of the associated leading edge genes (listed in Fig. 6B) revealed genes involved in ER stress response and protein folding/modification (Fig. 6C). Collectively, these data suggest that either T1D donors have fewer IFN-α responder cells to begin with or that these cells are eliminated selectively during disease development.

**Figure 6:**
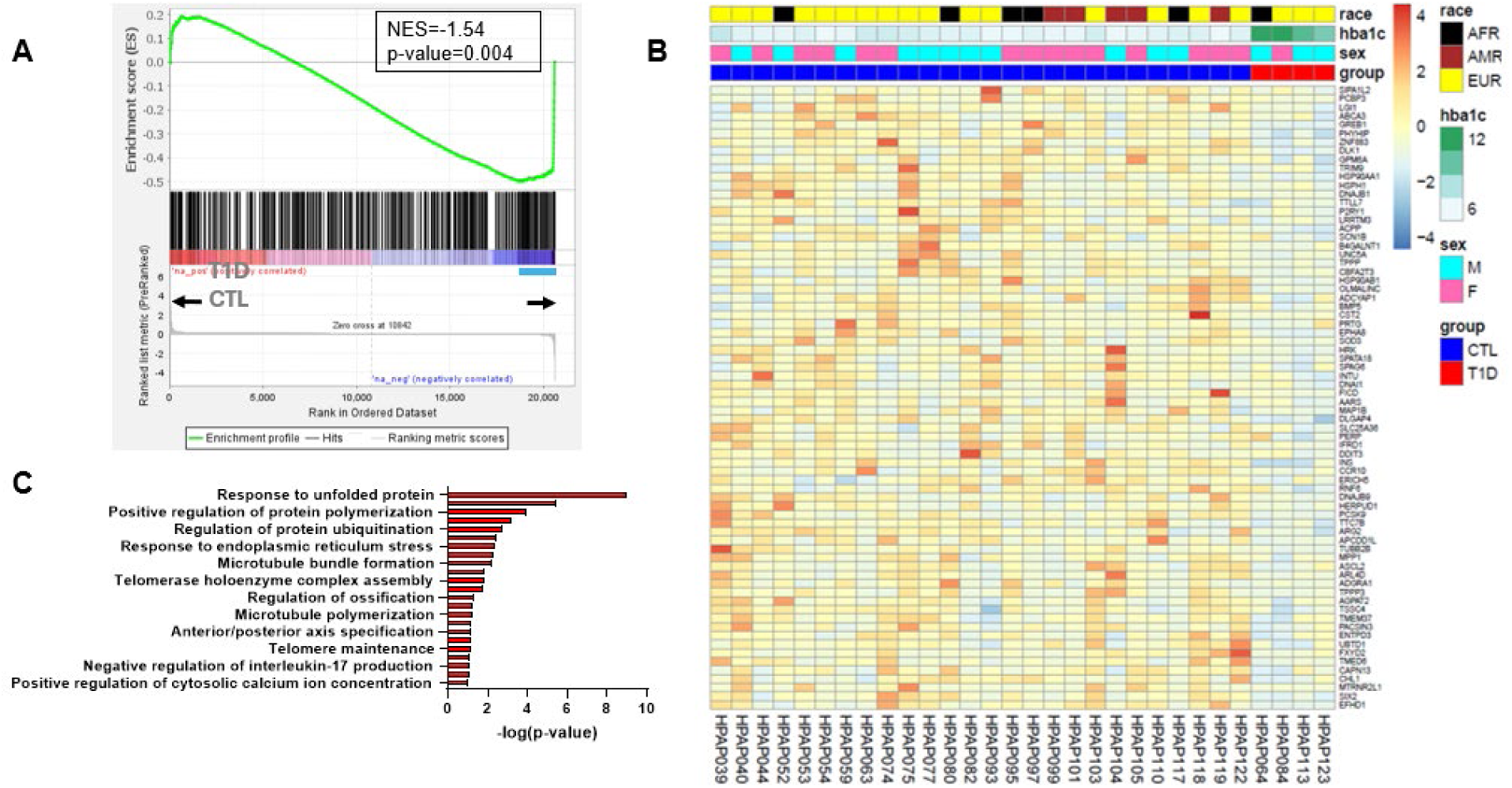
β-cell expression of responders’ genes in T1D and control HPAP donors. (A) GSEA enrichment plot for the up-regulated genes in responders across genes ranked by their shrunk log2FC of β-cell expression in T1D versus control donors from the HPAP scRNA-Seq dataset. The leading edge (72 genes) is indicated in cyan. Normalized Enrichment Score (NES) and nominal p-value are indicated. (B) Heatmap showing the DESEq2 normalized HPAP β-cell expression of these 72 leading edge genes. (C) Gene ontology analysis of these 72 leading edge genes. Values are represented as the –log(p-value) for each pathway.

### Human β-cells Accumulate Mitochondrial ROS *in vitro* Following IFN-α Treatment

To identify if the cytoplasmic ROS increase observed both *in vivo* and *in vitro* in response to IFN-α first originated in mitochondria, an organelle that is known for high amounts of superoxide production, we stained a human β-cell line (EndoC-βH1) with MitoSOX Green (Supp Fig 4). MitoSOX green is a dihydroethidium-based fluorescent dye that detects superoxide levels in mitochondria (35). We observed an increase in mitochondrial superoxide following one hour of IFN-α treatment (Supp Fig 4C). When pre-treating cells for six hours with 1μM MitoQ (36), a mitochondrially targeted antioxidant, this increase is abolished (Supp Fig 4C). To further investigate the timing of cytoplasmic and mitochondrial ROS production in human β-cells, we transfected EndoC-βH1 cells with a plasmid containing RIP1-GRX1-roGFP2 to measure cytosolic ROS as above, or TOMM20-GRX1-roGFP2, a mitochondrially targeted ROS biosensor. Two days following transfection, cells were treated with a vehicle, human IFN-α, human IFN-γ, or antimycin A. Images were collected from each RIP1-GRX1-roGFP2 dish at 15-minute (Fig 7A), 30-minute (Fig 7B), and 60-minute (Fig 7C) timepoints following addition of treatment. We found that at all these timepoints following addition of IFN-α there was a significant increase in cytoplasmic ROS production. Additionally, our studies with the TOMM20-GRX1-roGFP2 biosensor yielded similar results, with IFN-α stimulating a significant increase in ROS at the 15-minute (Fig 7D), 30-minute (Fig 7E), and 60-minute (Fig 7F) timepoints. These data demonstrate that the ROS produced in human β-cells in response to IFN-α likely originates within the mitochondria and then is rapidly released into the cytosol.

**Figure 7:**
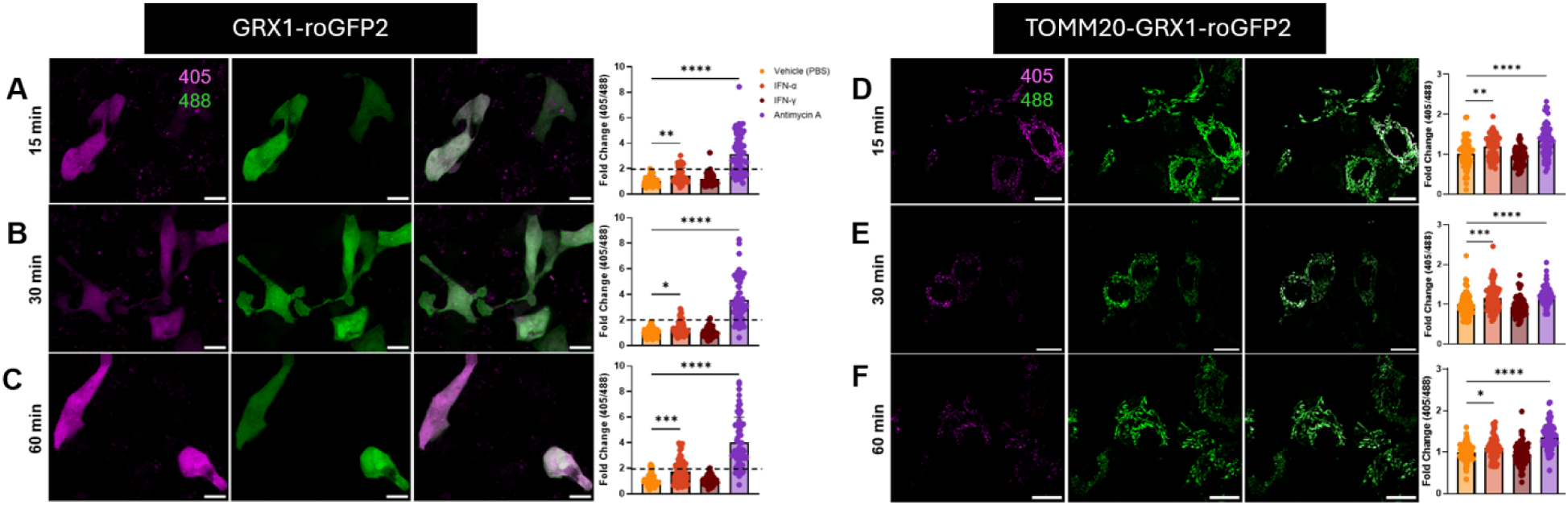
Human β-cells Accumulate Cytoplasmic and Mitochondrial ROS *in vitro* Following IFN-α Stimulation. (A-C) Representative images of EndoC-βH1 cells transfected with RIP1-GRX1-roGFP2 plasmid and treated with IFN-α for 15 min, 30 min, and 60 min, respectively. Scale bar length is 10μm. For analysis, the fold change of 405nm/488nm intensity was quantified from the average vehicle ratio. Single cell values from a total of three replicates were plotted and analyzed by one-way ANOVA, with variation plotted as mean ±SD. (D-F) Representative images of EndoC-βH1 cells transfected with TOMM20-GRX1-roGFP2 plasmid and treated with IFN-α for 15 min, 30 min, and 60 min, respectively. Scale bar length is 10μm. For analysis, the fold change of 405nm/488nm intensity was quantified from the average vehicle ratio. Single cell values from a total of three replicates were plotted and analyzed by one-way ANOVA, with variation plotted as mean ±SD.

## DISCUSSION

T1D is a complex autoimmune disease, with both a genetic predisposition and an unknown environmental triggering event, leading to the progressive loss of insulin-producing β-cells in the pancreatic islet. Prior literature has suggested that childhood viral infections may play a key role in driving T1D pathogenesis, possibly via ‘viral mimicry’, or similarity of viral epitopes to those presented by beta cells (37, 38). In fact, enteroviruses have been reported in the pancreata of T1D patients (39), although recent data do not support persistent infection of the pancreas as a common occurrence in T1D(40). Recently, insufficient editing of endogenous dsRNA molecules has been suggested as another possible trigger for the inflammatory response in autoimmune diseases, including T1D(41). Type 1 IFNs, including IFN-α, produced during the innate immune response to viral infection or in response to insufficiently edited endogenous RNAs, have long been implicated in T1D pathogenesis. Canonical interaction of type 1 IFNs with their receptors (IFNAR1, IFNAR2) activates the JAK/STAT pathway, where the transcription factors STAT1/STAT2 are phosphorylated and translocated to the nucleus (42). This translocation leads to stimulation of Interferon stimulated genes (ISGs) which function to mount a robust anti-viral response. Multiple case studies have reported the induction of insulin-dependent diabetes in patients prescribed type 1 IFNs as treatments for other medical conditions (43). Several studies have also demonstrated the presence of a strong type 1 IFN signature in the sera and tissues of individuals or donors with newly-onset T1D (3–5), as well as in the sera of children at risk for T1D development (1, 2). This has prompted interest in the use of small molecule inhibitors of the downstream IFN-α effectors such as tyrosine protein-kinase 2 (TYK2) as an approach to prevent T1D, with recent results demonstrating that they can elicit a protective effect in a mouse model of T1D (44). However, a critical gap in knowledge still exists as to how type 1 IFN signaling might facilitate autoimmune attack and β-cell dysfunction/destruction.

Prior studies have pointed towards IFN-α-induced islet HLA-I overexpression, making beta cells more ‘visible’ to the adaptive immune system, and ER stress as a mechanism by which IFN-α signaling could lead to autoimmune diabetes (6). Studies in other cell types have also shown that IFN-α induces ROS production (45–47). Although ROS have been implicated in detrimental conditions such as oxidative stress, other studies have identified ROS as key signaling molecules, promoting beneficial cellular processes such as β-cell proliferation and glucose-stimulated insulin secretion (11–13). ROS are mostly naturally generated as byproducts of the electron transport chain in the mitochondria, a process that is essential for glucose metabolism (48). As complex signaling molecules, ROS have also been identified as essential molecules in promoting the antiviral response (49–52), with several studies showing that mitochondrial ROS production promotes activation of the antiviral response.

Most islet biology studies, especially those utilizing human islets, have been performed in *in vitro* settings, removing islets from their niche. The technique of intravital microscopy allowed us to image islets in a pseudo-physiological condition following transplantation of human islets under the kidney capsule of immunodeficient mice (53). By coupling use of a β-cell specific redox biosensor (RIP1-Grx1-roGFP2) with this technique, we were able to monitor redox changes in human β-cells *in vivo* in real time. Using this approach, we demonstrate a rapid, heterogenous production of ROS in human β-cells *in vivo* following IFN-α stimulation. Specifically, we identify heterogeneous ROS accumulation in approximately 20% of β-cells on average across donors studied. However, the response was highly variable across individual donors, with some donors not having any responding cells, and other donors having as much as 63% responding cells.

To dive deeper into why this heterogeneity among donors occurs, we utilized the Human Islet Phenotyping Program (HIPP) provided to investigators in the IIDP. This analysis revealed a strong negative correlation between IFN-α-induced ROS production and HbA1C/BMI, suggesting that healthier donors have a higher number of β-cells that produce ROS in the presence of IFN-α. Additionally, GSEA of our sequencing datasets with T1D and control HPAP scRNA-Seq datasets indicated that genes up-regulated in responders are enriched in genes with higher expression in controls as compared to T1D β cells. Combined with the fact that ROS has been implicated in immune activation, this information leaves us with the working hypothesis that the increase in ROS production in the presence of acute IFN-α exposure is a protective response.

The increase in ROS production observed *in vivo* was recapitulated *in vitro* using fluorescent ROS probes DHE and DCFDA to measure superoxide and hydrogen peroxide production, respectively. However, it is important to note that DHE may not be used to exclusively measure superoxide radicals, as literature has shown that it is oxidized by other ROS (54). However, *in vitro*, we observe a significant increase in DHE in response to IFN-α stimulation, but not in DCFDA intensity, suggesting that superoxide is indeed the major ROS accumulating in human islets in response to IFN-α.

Since several studies have shown that specifically mitochondrial ROS promotes activation of the immune response (49, 51, 52), we asked if IFN-α induces mitochondrial ROS production in human β-cells. Indeed, we observed that IFN-α induces mitochondrial ROS production in human β-cells, and this ROS elevation is reversible with MitoQ pretreatment. Additionally, when comparing the gene signatures of non-responders and responders, we find that there is increased expression of several pro-survival and mitochondrial quality control genes coupled with a decrease in the gene expression of mitochondrial superoxide antioxidant, *SOD2*, supporting that there are differences in these genes that correlate with phenotypic stress response behavior.

In sum, we show that a subset of human β-cells produce ROS in response to IFN-α stimulation both *in vitro* and *in vivo*. This heterogenous increase is seen more robustly in human islet donors with lower BMI and HbA1C values, suggesting that it is a healthy response to cytokine stress. Further, when combined with analysis of the molecular signature of this subset of cells, our data suggests that prevalence of this cell subtype may be associated with more appropriate β-cell adaptive stress response that is predictive of cell survival. Interestingly, a recent meta-analysis and systematic review suggested that increased early life contact with microbes in children may be associated with a reduced risk of T1D (55). It has also long been known that the nonobese diabetic (NOD) mouse model of autoimmune diabetes exhibits a reduced incidence of disease when not housed in a pathogen free environment (56, 57). Collectively, these data are aligned with the hygiene hypothesis that has been historically attributed primarily to an immune cell-mediated effect (58, 59). Given the broad interest in the contribution of β-cells to T1D pathogenesis (54, 60–63), it will be imperative moving forward to determine the role that heterogeneity in the β-cell adaptive stress response plays in this process and how acute vs. chronic exposure to IFN-α alters the landscape of heterogeneity.

## MATERIALS/METHODS

### Sex as a Biological Variable

Sex was considered as a biological variable during the planning of all experiments involving both animals and human specimens. Both male and female mice were used for transplantation studies. Human islets used for experiments were accepted from both male and female donors, as indicated in Table 1.

### Study Approval

All animal experiments detailed in this manuscript were approved by the Indiana University School of Medicine Institutional Animal Care and Use Committee (IACUC). All human samples used in this study were de-identified specimens exempt from IRB approval.

### Human Islets

Human islets were obtained via the Integrated Islet Distribution Program (IIDP) or Alberta Diabetes Institute (ADI) islet core. Islets were cultured in non-tissue culture treated dishes containing 0.22μm filter sterilized standard Prodo islet media (Prodo CAT#PIM-S001GMP) supplemented with 5% human AB serum (Prodo CAT#PIM-ABS001GMP), 1% Glutamine and Glutathione (Prodo CAT#PIM-G001GMP), and 10mg/mL Ciprofloxacin (Fisher Scientific CAT#MT61277RG). Upon arrival, islets were centrifuged at 350g for 5 min to remove shipment media and then gently resuspended in complete islet media to recover in a dish overnight prior to experimentation.

### EndoC-βH1 Cell Culture

Human EndoC-βH1 cells were cultured in a monolayer and maintained on 100μg/mL Matrigel (Corning CAT#356237) and 2μg/mL fibronectin (Sigma-Aldrich CAT#F1141) precoated flasks and dishes. Cells were placed in 0.22μm filter sterilized low glucose DMEM media containing a final concentration of 2% albumin from bovine serum fraction V (Equitech-Bio CAT#BAH66), 10mM nicotinamide (Sigma-Aldrich CAT#N0636), 5.5 μg/mL transferrin (Sigma-Aldrich CAT#T8158), 6.7ng/mL sodium selenite (Sigma-Aldrich CAT#S5261), 50μM 2-mercaptoethanol (Sigma-Aldrich CAT#M3148), and Penicillin (100units/mL)/Streptomycin(100μg/mL) (Fisher Scientific CAT#15140122). Cells were split at ∼70-80% confluency, and media was changed every 4 days. Cells were cultured in incubators maintained at 37°C and 5% CO_2_.

### Mouse Husbandry

Mice were housed in a temperature-controlled facility with a 12 hr light/12 hr dark cycle and were given free access to standard food and water. For xenograft studies, male and female NOD.Cg-Prkdcscid Il2rgtm1Wjl/SzJ (NOD/SCIDγ(−/−)) (NSG) mice, aged 8-18 weeks, were used. These mice were obtained from the on-site breeding colony at the Preclinical Modeling and Therapeutics Core (PMTC) at Indiana University Simon Comprehensive Cancer Center at 6 weeks of age.

### Transduction of Islets

Human islets were obtained from islet resources as previously described. Following overnight recovery, islets were centrifuged at 350 g for 5 min at room temperature. Recovery media was removed, and the islet pellet was resuspended with PBS. The islets were centrifuged as previously described to form another pellet. The PBS supernatant was removed, and the islets were partially dissociated with Accutase (Innovative Cell Technologies CAT#AT-104) for 1 minute at 37°C. The Accutase enzyme was inactivated with complete islet media and removed following centrifugation as previously described. Islets were then transduced with 1*10^11^ PFU of adenoviral RIP1-Grx1-roGFP2 per 1000 IEQs for ∼16-18hrs at 37°C in complete islet media.

### Islet Transplantation

Following transduction, islets were centrifuged at 350g for 5 min, and viral islet media was removed. Islets were transferred to 100μL of complete human islet media for transplantation under the left kidney capsule of NSG mice by the Indiana University Diabetes Center Islet Core and in house. Approximately 1,000 IEQs were transplanted per mouse for xenograft studies. 4 000 IEQs were obtained per donor, with transplantation into four separate mice conducted to account for variability. Briefly, transplant recipients were subcutaneously injected with buprenorphine at a dose of 3.25mg/kg prior to surgery. Mice were then anesthetized by continuous 2.5% isoflurane inhalation and placed atop a warming blanket on which they remained until reflexes were regained post-surgery. Throughout the surgery, ophthalmic ointment (Artificial Tears gel) was directly applied to the mouse’s eyes to avoid drying. The left flank region was shaved, and any remaining hair was depilated with a hair removal cream prior to sterilization with ethanol swab. A small incision was made in the skin above the left kidney. The abdominal wall was gently incised exposing the kidney. The left kidney was gently externalized with cotton swabs and moistened with saline. The kidney was continuously administered saline during this procedure to avoid drying and tearing of the renal capsule. Islets were placed in sterile polyethylene (PE-50) tubing that was distally attached to a Hamilton screw syringe equipped a blunt needle tip. Tubing containing islets was beveled with surgical scissors. A small nick was made on the renal capsule via a 30G needle, and the beveled end of tubing was carefully placed under the kidney capsule. The islets were slowly delivered under the capsule using the Hamilton syringe. The kidney was gently placed back into its location in the peritoneal cavity, and the incisions were non-continuously sutured using absorbable 5-0 vicryl filament. Post-surgery, the animals were rehydrated with saline and administered analgesics (1.3mg/mL extended-release buprenorphine) for pain relief. Animals were monitored until they regained reflexes. Throughout the surgery, the animal’s temperature was maintained between 35°C-36.7°C. Following surgery, animals were singly housed in a clean cage and provided with wet feed.

### Intravital Imaging

Two weeks post-islet transplant, animals were anesthetized with 2.5-3% isoflurane circuit and kept under anesthesia throughout the imaging session. Animals were placed on a heating pad, and ophthalmic ointment was applied to both eyes. The left flank of the animal was depilated and sterilized with an ethanol swab. A small incision was performed along the post-transplant sutures to expose the left kidney. The kidney was gently externalized with cotton swabs and placed under a 40mm coverslip glass bottom dish. The animals were gently flipped in an orientation so that the kidney was beneath the animal, pressed against the coverslip for subsequent imaging. Islets were identified using GFP fluorescence through the eyepiece. Islets were imaged using either a Leica SP8 or Leica SP8 DIVE microscope equipped with a HCX IR APO 25x/0.95 W lens. During confocal imaging, RIP1-Grx1-roGFP2 was sequentially excited by 405nm/488nm laser stimulation, and emission was collected between 490 and 600nm using internal hybrid detectors as described previously (29). Following baseline acquisition, mice were injected with saline or 2.75×10^5^ IU/mL of recombinant, carrier-free human IFN-α (Novus Biologicals CAT#NBP2-26551) via retro-orbital injection. Z-stack images (30-50μm total thickness, 5μm step size) were collected every 2-3 min for 30-60 min. Following imaging of treatment conditions, animals were injected with a vasculature dye (AlexaFluor647-conjugated Albumin, produced in house) to visualize blood flow, and a final image was collected with excitation at 650nm and collection at 670-700nm. Animals were euthanized via cervical dislocation. Terminal blood was collected via cardiac puncture, and tissues were harvested for downstream analysis.

### Quantification of *in vivo* Redox Status of β-cells

Raw images from the intravital experiment were first processed using rolling ball background subtraction (radius of 50 pixels), and a Gaussian blur filter was applied. Max intensity projections were created for each islet analyzed. A 3D drift correction (Image Stabilizer plugin) was applied to datasets affected by motion artifacts caused by the animal’s breathing. Regions of interest (ROI) were drawn manually around each cell expressing sensor using the 488nm channel. Mean fluorescence intensities (405nm and 488nm) were calculated for each ROI for every timepoint using ImageJ (64). The ratio of fluorescence intensity (405nm/488nm) was calculated for each ROI to determine oxidative state. Additionally, fold change in ratio, measured by ratio change from baseline, was calculated for each ROI to determine changes in oxidative state. We identified responders as β-cells that had a fold change in 405nm/488nm ratio of 2 or higher when compared to baseline.

### Functional Analysis of Transplanted Islets

Following imaging, terminal blood was collected and centrifuged at 21,000 g for 5 min at room temperature. Sera was transferred to a new centrifuge tube and stored at −80°C. Sera was used to assess human C-peptide levels to determine the functionality of transplanted human islets. An ALPCO Insulin ELISA kit was used for this calculation, and all assessments were performed by the Indiana University Center for Diabetes and Metabolic Diseases Translational Core. For analysis of serum IFN-α levels, mesoscale discovery (MSD) analysis of human IFN-α was conducted at the Indiana University Center for Diabetes and Metabolic Diseases Translational Core, according to manufacturer’s instructions.

### Tissue Embedding

Tissues collected after intravital imaging of human islets (kidneys containing transplanted islets) were fixed with 4% PFA and embedded in paraffin blocks for downstream immunofluorescence analysis.

### Immunofluorescence Staining

Tissues containing embedded islets were sectioned at 5μm thickness and stained by immunofluorescence using the following primary antibodies: anti-pSTAT2 (Cell Signaling Technology, CAT#88410, 1:200), anti-insulin (Dako, CAT#IR002, 1:10), anti-4-HNE (Abcam, CAT#ab48506, 1:100), and anti-8OHdG (Abcam, CAT#ab62623, 1:100). For secondary antibodies, Alexa Fluor 647 donkey anti-rabbit IgG (Invitrogen, CAT# A31573), Alexa Fluor 594 goat anti-mouse IgG (Invitrogen, CAT# A11032), and Alexa Fluor 488 goat anti-guinea pig (Invitrogen, CAT# A11073) IgG antibodies were used. Tissues were stained with DAPI to mark nuclei. Images were acquired using a Zeiss LSM 800 confocal microscope or a Leica Stellaris confocal microscope.

### Live Imaging of Human Islets with ROS Probes

Human islets were obtained from islet resources as previously described. Following overnight recovery, roughly 500 IEQs were transferred into 2mL of fresh complete human islet media in 6-well plates for addition of treatment conditions. Following one-hour treatment with PBS vehicle, 2000IU/mL recombinant, carrier-free human IFN-α (Novus Biologicals CAT#NBP2-26551), 10μM Antimycin A (Sigma-Aldrich CAT#A8674) or 20μM hydrogen peroxide (Sigma-Aldrich CAT#216763), islets were transferred into new wells containing 2μM Hoechst (Thermo Scientific CAT#62249) and 10μΜ dihydroethidium (Cayman Chemical CAT#12013) or 10μΜ 2’,7’-dichlorodihydrofluorescein diacetate (Cayman Chemical CAT#20656) in 2mL complete human islet media for 25 minutes. The islets were then transferred to an imaging dish containing fresh complete human islet media just prior to imaging. Images were collected on a Zeiss LSM800 confocal microscope or a Leica Stellaris confocal microscope. Z-stack images of at least 3 whole islets per condition were collected. Hoechst was imaged via excitation at 405nm and emission collection at 413-491nm. Dihydroethidium was collected via excitation at 488nm and emission collection at 590-640nm. 2’,7’-dichlorodihydrofluorescein diacetate was collected via excitation at 488nm and emission collection at 498-610nm.

### Preparing Human Islets for RNA Sequencing

Human islets were obtained from islet resources as previously described. Following overnight recovery, islets were divided and treated for one hour with PBS vehicle (2000 IEQ), 2000 IU/mL recombinant, carrier-free human IFN-α (Novus Biologicals CAT#NBP2-26551) (2000 IEQ), or 10μM Antimycin A (Sigma-Aldrich CAT#A8674) (500 IEQs). A subset of islets remained untreated for a negative control (500 IEQ). Following treatment, islets were pelleted at 350g for 5 min and dissociated in Accutase (Innovative Cell Technologies CAT#AT-104) for 15 minutes in a 37°C water bath under gentle agitation. Following dissociation, single cells were filtered through a 30μM mesh filter, and the cell suspension was transferred on ice to the fluorescence-activated cell sorting (FACS) instrument. A blue laser equipped with a PE filter was used for the sorting experiments on the FACS instrument. The unstained, untreated negative control population was run first to gate true negative cells. Five minutes prior to running the Antimycin A sample, 10μΜ dihydroethidium (Cayman Chemical CAT#12013) was added to the tube and left to incubate at room temperature in the dark. After the Antimycin A sample was run to gate the positive population of cells, the vehicle and IFN-α treated samples were each stained with 10μΜ dihydroethidium and sorted based off the presence or absence of reactive oxygen species (ROS) as determined by the established gates. RNA from each sample was isolated using the Qiagen RNeasy Mini Kit (CAT#74134) protocol with addition of DNase (Qiagen CAT#79254).

### Bulk RNA Sequencing of Human Islets

Sequencing analysis was carried out in the Center for Medical Genomics at Indiana University School of Medicine, which is partially supported by the Indiana University Grand Challenges Precision Health Initiative. Paired analysis was conducted for each human islet donor. After passing quality control checks, 1ng of RNA from all samples were run on an Illumina NovaSeq X Plus sequencer. Library prep was conducted using the Clonetech SMARTseq V4 kit. Raw sequencing data was processed and analyzed for differential gene expression by the Medical Genomics Core using EdgeR, and reads were aligned to the hg38 genome using STAR aligner. Low quality mapped reads (including reads mapped to multiple positions) were excluded, resulting in analysis of 17,192 genes. Differentially expressed genes were identified following an FDR correction to have an adjusted p-value of less than 0.05.

### Comparison of Datasets

To assess the β-cell expression in T1D versus control donors for the genes up-regulated in the responder population, we leveraged the publicly available scRNA-Seq data from the Human Pancreas Analysis Program (HPAP). For each donor islet sample, we obtained counts with CellRanger v7.1.0(65) and utilized Seurat v4.9.9.9041(66) (complemented by SoupX(67) and scDblFinder(68)) for clean-up, normalization, pre-processing and data integration. Cells were annotated with two approaches: scSorter(69) and manual cluster annotation based on selected pancreatic cell markers. Only cells for which the two approaches agreed were assigned to a final cell type. We then used HPAP samples having a minimum number of 100 β-cells and whose libraries had been prepared with the same chemistry kit (SC3Pv3). For each donor, β-cell pseudo-bulk counts were generated and 20,621 genes with ≥10 counts in ≥ 4 HPAP donors were retained for analyses. DESeq2(70) was used to compare the available 4 T1D to 26 control donors, generating a pre-ranked gene list based on shrunk log2 fold change which was then input into a GSEA analysis(34). GSEA was used to study enrichment of the gene set of up-regulated genes in responders (441 genes with adjusted p-value<0.05 and FC > 1.5). and revealed that these genes were enriched in genes with higher β-cell expression in control donors. The 72 leading edge genes from this analysis were further examined with Gene Ontology analysis.

### Gene Ontology (GO) Analysis

Genes of interest were first identified via a p-value less than 0.05 and a logFC of 1.5. The official gene symbol of these identified genes was analyzed using the NIH Database for Annotation, Visualization and Integrated Discovery (DAVID) program. Briefly, the Functional Annotation Tool was utilized to conduct GO analysis of identified genes. These pathways were subsequently analyzed to be represented as a –log(p-value).

### Live Imaging with EndoC βH1 Cells

For imaging preparation, EndoC-βH1 cells were split into imaging dishes (MatTek CAT#P35G-1.5-14-C) at a seeding density of roughly 100,000 cells/dish. Imaging experiments were conducted 1-3 days following cell seeding. A subset of dishes was pre-treated with 1μM of MitoQ (Selleck CAT#S8978) for six hours. Cells were treated for one hour with PBS vehicle, 2000IU/mL recombinant, carrier-free human IFN-α (Novus Biologicals CAT#NBP2-26551), or 10μM Antimycin A (Sigma-Aldrich CAT#A8674). Treatment media was removed and replaced with 2mL of fresh, complete media containing 2μM Hoechst (Thermo Scientific CAT#62249), 0.5μM MitoSOX Green (Invitrogen CAT#M36006), and 0.1μM MitoTracker Deep Red (Invitrogen CAT#M22426) for 25 minutes. Fluorescent probe media was removed and replaced with 2mL of fresh, complete media for imaging at a Leica Stellaris confocal microscope. Z-stack images of several areas on the dish per condition were collected. Hoechst was imaged via excitation at 405nm and emission collection at 413-491nm. MitoSOX Green was collected via excitation at 488nm and emission collection at 498-610nm. MitoTracker Deep Red was collected via excitation at 638nm and emission collection at 646-811nm.

For biosensor imaging studies, TOMM20-roGFP2 and RIP1-GRX1-roGFP2 plasmids were utilized. EndoC-βH1 cells were split into imaging dishes (MatTek CAT#P35G-1.5-14-C) at a seeding density of roughly 150,000 cells/dish. One day following seeding, cells were either transfected with 3μg of TOMM20-GRX1-roGFP2 or RIP1-GRX1-roGFP2 plasmid using the Lipofectamine® 2000 protocol. Roughly 36 hours post-transfection, cells were treated for one hour with PBS vehicle, 2000IU/mL recombinant, carrier-free human IFN-α (Novus Biologicals CAT#NBP2-26551), 1000 IU/mL recombinant, carrier-free human IFN-γ (R&D Systems CAT#285-100/CF), or 10μM Antimycin A (Sigma-Aldrich CAT#A8674). Z-stack images were collected at 3 areas of each dish at 15 min, 30 min, and 60 min timepoints using a Leica Stellaris confocal microscope. The dishes were subsequently excited at 405nm and 488nm, with emission collection set between 498-610nm. For these experiments, the fold change in the RIP1-GRX1-roGFP2 ratio (405nm/488nm) was calculated from the average vehicle condition for each timepoint. For analysis, a one-way ANOVA was conducted, with results plotted as mean ± SD.

### Statistical Analysis

Images were analyzed using Fiji ImageJ software. Groups were compared by one-sample t-test or one-way ANOVA, using GraphPad Prism, and the results are represented as mean±SD. P-values < 0.05 were considered statistically significant. For correlation analysis, data from each donor was recorded, and linear regression analysis was conducted in GraphPad Prism.

### Data Availability

The Supporting Data Values file will be made available to the public upon acceptance of this manuscript. RNA sequencing datasets will be made available to the public via GEO. The HPAP scRNA-Seq data and meta-data are available at the PancDB website (https://hpap.pmacs.upenn.edu/).

## CONTRIBUTIONS

LEW and AKL were responsible for designing the research studies. LEW was primarily responsible for conducting experiments. LEW, OM, MMMI, MCA, CM, and BED conducted intravital imaging experiments. KSO, BED, and OM conducted human islet transplantations. LEW, EM, and OM conducted data analysis. LEW, EM, KHK, and AKL were responsible for writing the manuscript. All authors contributed to manuscript edits and approved of the submitted version.

## ACKNOWLEDGEMENTS

This work was made possible with the following grant funding: a new investigator award from NIDDK/HIRN RRID: SCR_014393 to AKL, a NIH/NIDDK New Investigator Gateway Award R03 DK127766 to AKL, NIH/NIDDK grant R01DK124380 to AKL, and NIH/NIDDK grant F31DK137567 to LEW. Additional support provided by the Herman B Wells Center for Pediatric Research was in part from the Riley Children’s Foundation. The authors acknowledge the support of the Microscopy Core and Islet and Physiology Core of the Indiana Diabetes Research Center (P30DK097512). The authors thank the members of the Indiana University Melvin and Bren Simon Comprehensive Cancer Center Flow Cytometry Core for their outstanding technical support. Human pancreatic islets and/or other resources were in part provided by the NIDDK-funded Integrated Islet Distribution Program (IIDP) (RRID:SCR_014387) at City of Hope, NIH Grant # U24DK098085. Human islets for research were in part provided by the Alberta Diabetes Institute IsletCore at the University of Alberta in Edmonton (http://www.bcell.org/adi-isletcore.html) with the assistance of the Human Organ Procurement and Exchange (HOPE) program, Trillium Gift of Life Network (TGLN), and other Canadian organ procurement organizations. This manuscript used data acquired from the Human Pancreas Analysis Program (HPAP-RRID:SCR_016202) Database (https://hpap.pmacs.upenn.edu/), a Human Islet Research Network (RRID:SCR_014393) consortium (UC4-DK-112217, U01-DK-123594, UC4-DK-112232, and U01-DK-123716). Islet isolation was approved by the Human Research Ethics Board at the University of Alberta (Pro00013094). All donors’ families gave informed consent for the use of pancreatic tissue in research. We would like to thank the organ donors and their families for making this work possible.

---

**Supplementary Figure 1:**
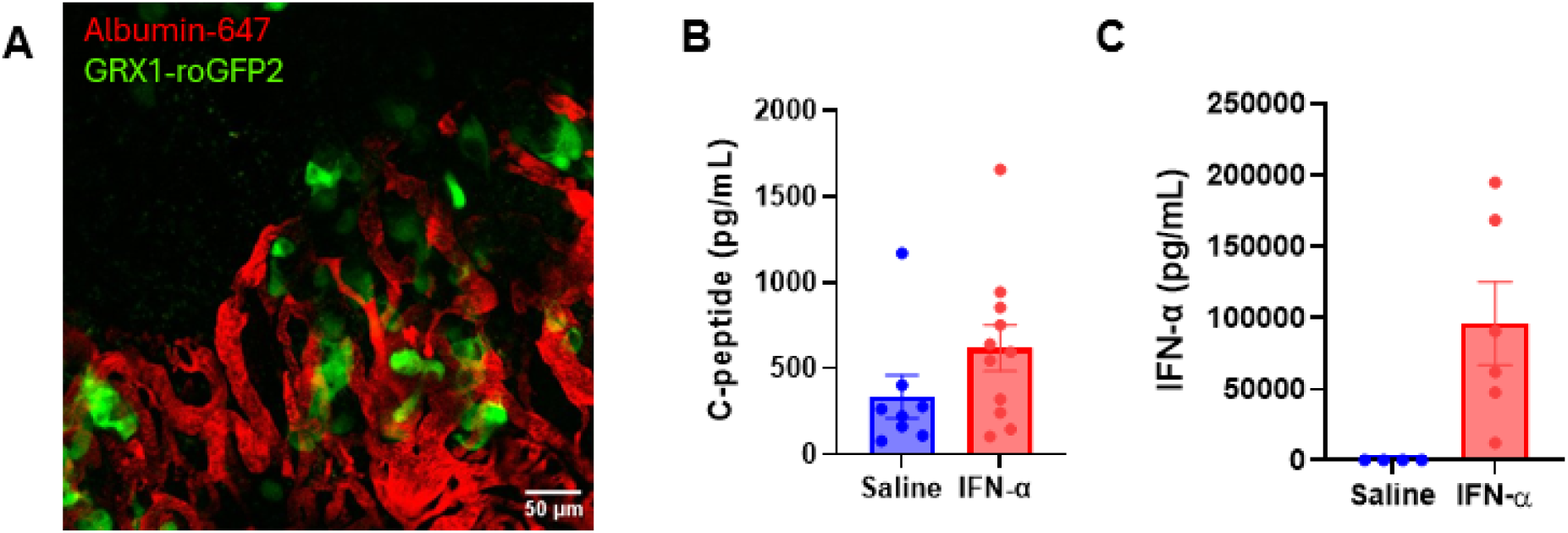
Transplanted Islets are Well-Vascularized and Functional. (A) Representative image of an islet expressing roGFP2 (green) surrounded by vasculature marked by Albumin-AlexaFluor647 (red). (B) Quantification of human C-peptide levels in the sera of recipient mice that received human islet transplants under the kidney capsule. Saline injected mice are displayed in blue, and IFN-α injected mice are shown in red. (C) Mice with transplanted islets were retro-orbitally injected with IFN-α as conducted during intravital imaging. IFN-α levels were quantified in both saline (blue) and IFN-α (red) injected mice.

**Supplementary Table 1:** Individual Intravital Imaging Donors Percent Responders. The number and percentage of responders for each donor are plotted for the 30 min, 45 min, and 60 min timepoints. Both saline and IFN-α stimulated islet cell percentage count is displayed.

| Donor ID | # Responders 30 Min. Saline | # Responders 45 Min. Saline | # Responders 60 Min. Saline | # Responders 30 Min. IFN-α | # Responders 45 Min. IFN-α | # Responders 60 Min. IFN-α |
| --- | --- | --- | --- | --- | --- | --- |
| SAMN17367763 | 0/57 (0%) | 0/57 (0%) | 0/57 (0%) | 2/52 (3.8%) | 1/52 (1.9%) | 5/52 (9.6%) |
| SAMN16423389 | 1/61 (1.6%) | 2/61 (3.2%) | 0/61 (0%) | 16/45 (35.6%) | 16/45 (35.6%) | 16/45 (35.5%) |
| SAMN16427178 | 0/42 (0%) | 0/42 (0%) | 0/42 (0%) | 0/13 (0%) | 0/13 (0%) | 0/13 (0%) |
| SAMN28450743 | 0/20 (0%) | 0/20 (0%) | 0/20 (0%) | 20/66 (30.3%) | 20/66 (30.3%) | 14/66 (21.2%) |
| SAMN10920290 | Not imaged | Not imaged | Not imaged | 26/41 (63.4%) | 25/41 (61%) | Not imaged |
| R309 | Not imaged | Not imaged | Not imaged | 28/250 (11.2%) | Not imaged | Not imaged |
| R395 | 0/140 (0%) | 4/140 (2.9%) | 7/140 (5%) | Not imaged | Not imaged | Not imaged |
| R448 | 0/43 (0%) | 0/43 (0%) | 0/43 (0%) | 0/53 (0%) | 0/53 (0%) | 0/53 (0%) |

**Supplementary Figure 2:**
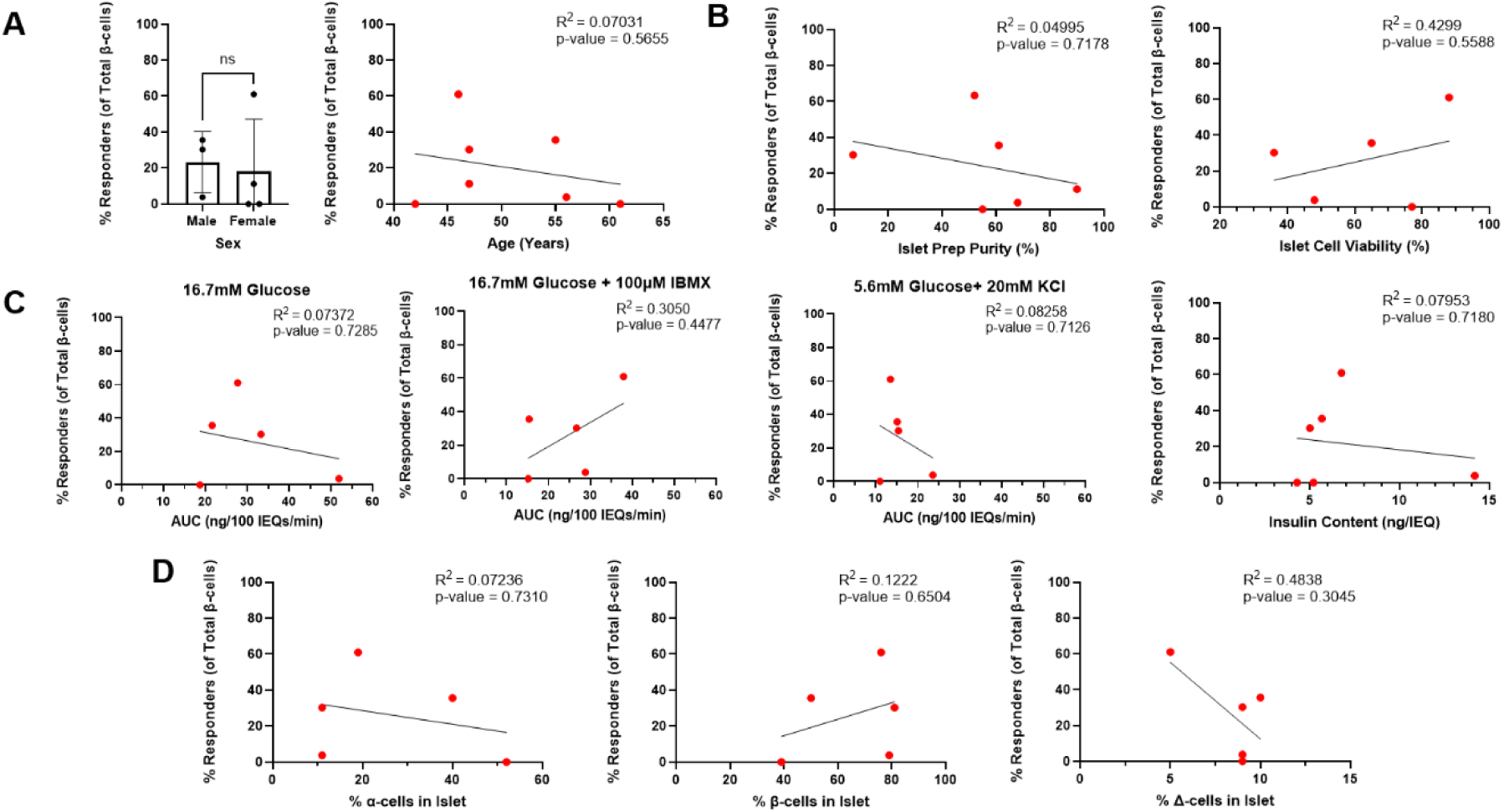
Human Islet Phenotyping Variables vs Percent Responders. (A) Demographic variables from donors, including sex and age, are plotted against percent responders of total β-cells at 30 min. (B) Islet preparation purity and viability are plotted against percent responders of total β-cells at 30 min. (C) Perifusion data from donors are plotted against percent responders of total β-cells at 30 min. (D) Islet cellular composition is plotted against percent responders of total β-cells at 30 min.

**Supplementary Figure 3:**
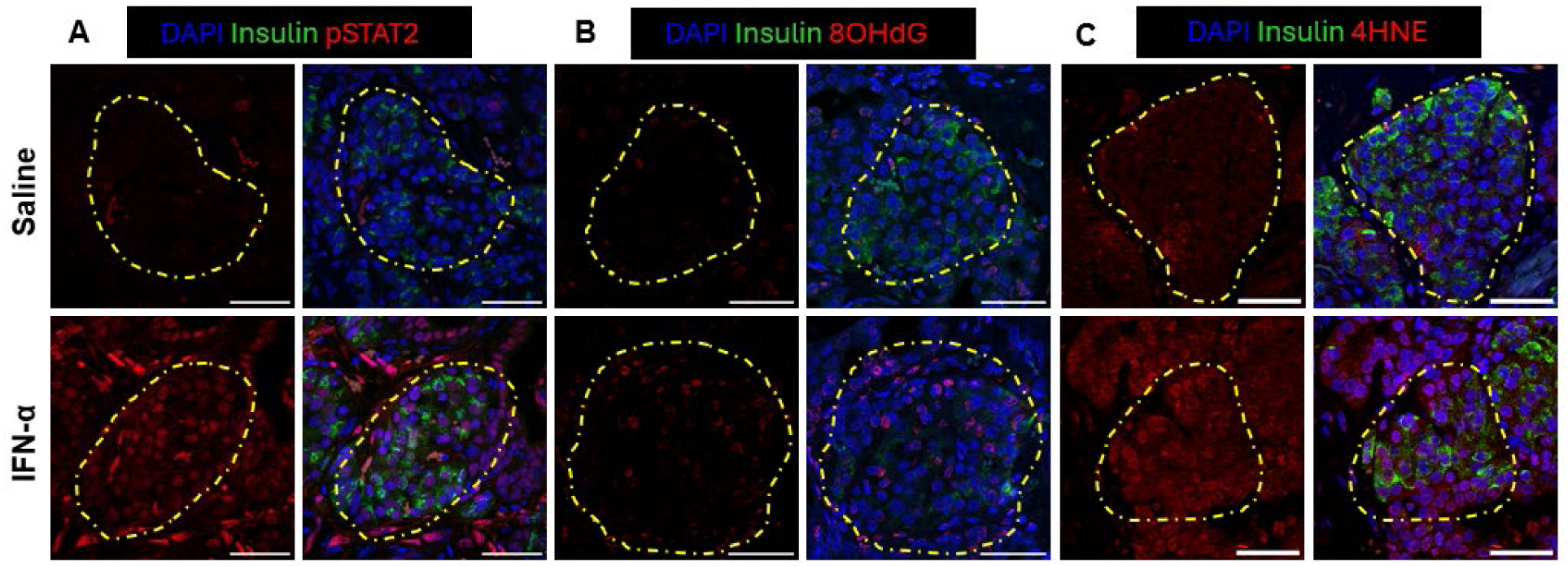
IFN-α Treatment Induces Acute Protein Changes in IFN-α Signaling and ROS Production. Representative images of immunofluorescence staining conducted on transplanted human islets. Saline stimulated islets are shown in the top row, and IFN-α stimulated islets are shown in the bottom row. Scale bar length is 50μm. (A) Islets stained for DAPI (blue), insulin (green) and pSTAT2 (red). (B) Islets stained for DAPI (blue), insulin (green), and 8OHdG (red). (C) Islets stained for DAPI (blue), insulin (green), and 4-HNE (red).

**Supplementary Figure 4:**
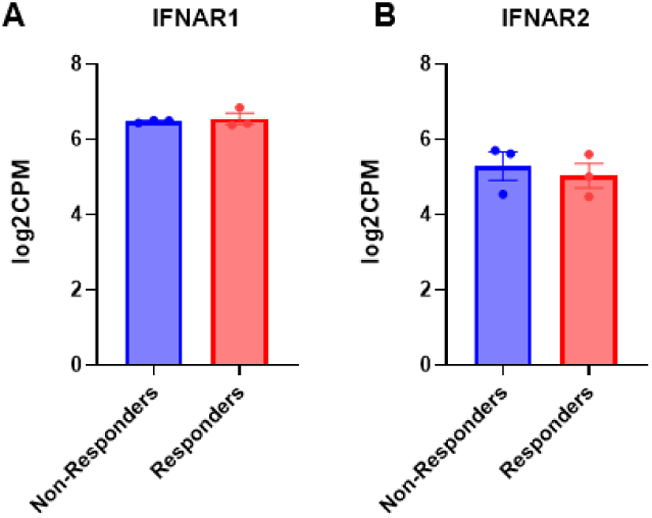
IFN-α Receptor Transcript Expression is not Different Between Responding and Non-Responding β-Cells. (A) Relative gene expression of non-responding (blue) and responding (red) islet cells from each donor used for RNA sequencing, represented at the log2cpm for the *IFNAR1* gene. (B) Relative gene expression of non-responding (blue) and responding (red) islet cells from each donor used for RNA sequencing, represented at the log2cpm for the *IFNAR2* gene. Analysis was conducted as a t-test, comparing the means of both treatments.

**Supplementary Figure 5:**
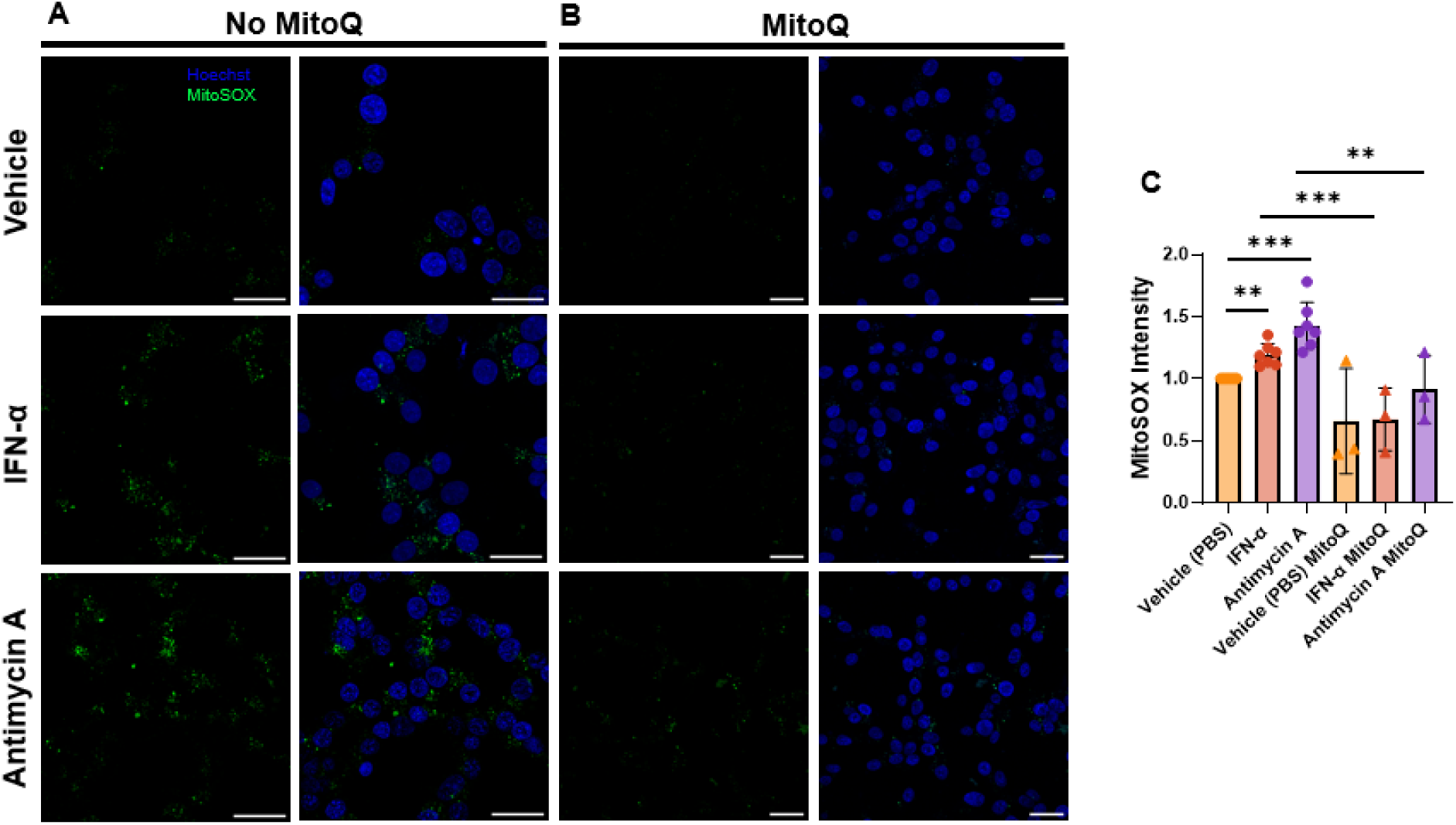
IFN-α Treatment Induces Mitochondrial ROS Production that is Reversible with MitoQ Pretreatment. Representative images of EndoC-βH1 cells stained with Hoechst (blue) and MitoSOX Green (green). Cells treated with PBS vehicle are displayed in the top row, cells treated with 2000IU/mL human IFN-α are displayed in the middle row, and cells treated with 10μM Antimycin A are shown in the bottom row. (B) Representative images of EndoC-βH1 cells pretreated with 1μM MitoQ for six hours stained with Hoechst (blue) and MitoSOX Green (green). (C) Quantification of MitoSOX Green mean fluorescence intensity in EndoC-βH1 cells. Analysis was conducted as a one-sample t-test, with data normalized to the vehicle without MitoQ pretreatment condition.

